# Neural signatures of prioritization and facilitation in retrieving repeated items in Visual Working Memory

**DOI:** 10.1101/2023.08.18.553911

**Authors:** Abhishek Singh Narvaria, Arpan Banerjee, Dipanjan Roy

## Abstract

Visual Working memory (VWM) is a limited-capacity system where working memory items compete for retrieval. Some items are maintained in the working memory in the “region of direct access,” which holds information readily available for processing, while other items are in a passive or activated long-term memory state and require cognitive control. Moreover, their recognition requires moving from the most active template in VWM to another one with the shift of attention. Stimulus properties based on similarity can link items together, which can facilitate their retrieval due to prioritization. To investigate the neural dynamics of differential processing of repeated versus not-repeated items in working memory, we designed a modified Sternberg task for testing recognition in a VWM-based EEG study where human participants respond to a probe for an item’s presence or absence in the representation of an encoded memory array containing repeated and not repeated items. Significantly slower response times and comparatively poor accuracy for recognising not-repeated items suggest that they are not prioritized. We identified specific differences in spectral perturbations for sensor clusters in the power of different frequency bands as the neural correlate of probe matching for not-repeated vs. repeated conditions, reflecting biased access to VWM items. For not-repeated item probe matching, delay in beta desynchronization suggests poor memory-guided action selection behaviour. An increase in frontal theta and parietal alpha power demonstrated a demand for stronger cognitive control for retrieving items for not-repeated probe matching by shielding them from distracting repeated items. In summary, our study provides crucial empirical evidence of facilitation and prioritization of repeated items over non-repeated items and explains the probable role of different EEG rhythms in facilitated recognition of repeated items over goal-relevant, not-repeated items in VWM.

## 1. Introduction

Working memory refers to our innate cognitive ability to temporarily hold and manipulate relevant information “in mind.” Two key features of working memory are its flexibility and its starkly limited capacity to maintain only a limited number of items (Adam, K. C., & Serences, J. T., 2019). In working memory (WM), items are held as mental representations that compete for retrieval, especially during probe comparison tasks. This competition for retrieval can impact how well we remember these items. For example, in a working memory recognition task, non-target items similar to the target create competition and interference. This competition can affect the speed and accuracy of retrieval (Olivers et al., 2011). In such tasks, an attentional template of the probe is created and is compared with different WM representations one after the other in a serial manner and requires attention to bring one representation at a time into an active state in working memory. This process continues till the item matching is completed. As per embedded processing models of working memory (Cowan, 1999; Oberauer, 2002, 2005), the items in WM are maintained in focus of attention as in the active state, while less relevant items are in the activated region of long-term memory and are considered to be in a passive state (LaRocque et al., 2014; Peters et al., 2019; Li et al., 2020, 2021; Zhang et al., 2022). These states determine how easily accessible these WM representations are during a memory retrieval task.

Prioritization and facilitation of items ease out this competition for retrieval. Items in the form of memorized arrays might lead to enhanced facilitation of WM representations due to perceptual features like their size or hue (Constant, M., & Liesefeld, H. R., 2021). This can be due to different mechanisms, which are mainly bottom-up. Regarding multi-item WM retrieval, shared inter-item properties like similarity or repetition of items (Guofang Ren et al., 2023; Hamblin-Frohman et al., 2023; Lin & Luck, 2009) can be facilitated. These inter-item properties can lead to repetition facilitation, where these items are bound together in the form of chunks, leading to enhancement in retrieval (Chekaf et al., 2016; Thalmann et al., 2019).

However, this item association is not always facilitatory and can lead to repetition inhibition, as this pattern of similarity or repetition needs to be detected and should not have any lag or be presented very distant from each other (Catherine L. Lee 1976; Robert G. Crowder. This way, WM items do not undergo Gestalt perception (Dwight J. Peterson et al., 2013). The failure to detect repetition leads to even inhibition as shown in the Ranschburg effect in short-term memory (Greene, R.L.,1991). However, if the pattern is detected, the items bound are chunked, and associative-linked memory items are automatically activated (Oberauer and Lange, 2009). The representation of identical items might lead to a lower activation threshold for their probe matching compared to non-facilitated items (Guofang Ren et al., 2023). Hence, these mechanisms are important to answer the question whether repetition of items in a working memory array can lead to better recognition and facilitation of their representations in a probe comparison task, and leads to conflict for non-facilitated items that are not repeated.

In line with the above, one testable hypothesis could be that repeated items (Rep) might be facilitated and interfere with probe matching for not-repeated items (NRep) in VWM, impacting their retrieval during working memory performance. Moreover, the competition for facilitated retrieval might depend on the role of attention in inhibiting irrelevant representations and response preparation in VWM. The probe matching for VWM representations must be reflected in the EEG data based on the alteration in brain oscillations and may shed crucial insight into their possible neural correlates for prioritization, facilitation, or hindrance in the VWM recognition task.

The probe matching window is important because it allows researchers to isolate the neural activity associated with matching a new input (the probe) with stored information in visual working memory (VWM). Different frequency bands, like alpha, theta, and beta, reflect different cognitive processes involved in this matching, including allocation of attention, memory maintenance, and conflict in decision making, The changes in frequency bands during the probe matching window provide critical insights into how attention is allocated to the matching probe, which is required for matching the attentional template to maintained VWM representations. During VWM, the dynamics of neural beta (13-30 Hz), especially in the central electrode, are associated with memory-guided behavior as it can affect motor preparation, as shown by previous studies (Boettcher et al., 2021; Nasrawi et al., 2023; Nasrawi & van Ede, 2022; Schneider et al., 2017). Such motor preparation signals index the access to items to be prioritized within VWM (Ding et al., 2024). Extant literature further suggests that the Alpha frequency band plays a crucial role in sensory information-specific attention requirements. Interestingly, changes in the relative priority of stored representations and maintenance are reflected in the modulation of posterior alpha power oscillations (8–14 Hz) as reflected in alpha lateralization for the selection of task-relevant information for the internal selection of information maintained within VWM (van Ede, F., Niklaus, M., & Nobre, A. C., 2017). Alpha desynchronization causes active inhibition of items that need to be inhibited. It can also inform about the matching of the attentional template of the probe with the relevant or irrelevant item’s representation.

Complex behavioral tasks require a higher need for cognitive effort and conflict monitoring (Cavanagh & Frank, 2014; Cavanagh & Shackman, 2015; Cavanagh et al., 2012. In line with this, previous studies have reported Theta power (4-7Hz) oscillatory changes related to top-down control over items (Sauseng, P., Griesmayr, B., Freunberger, 2010; Sauseng & Liesefeld, 2020). More specifically, it has been shown that the frontal theta controls the endogenous attentional selection mechanisms of task-relevant items (Johnson et al., 2017; Sauseng, P., Griesmayr, B., Freunberger, R., & Klimesch, W., 2010). Alteration in the power of Fronto-medial theta can resolve the conflict in probe matching for irrelevant items in the probe conditions.

In this study, to capture the bias in probe matching for repeated vs. not-repeated items in the VWM task and its neural correlates, we have conducted an EEG study that utilizes a memory array facilitating the encoding of items of both the repeated and not-repeated categories. The encoding of items of both categories with an equal chance of the appearance of a relevant probe. This is to gain empirical evidence for how WM representation of certain WM items is prioritized and behaviorally influences facilitation for repeated or not-repeated items as captured by individual response time and accuracy for responding to relevant probes. Next, we investigate their neural correlates using EEG to test our hypothesis that band-specific spectral power differences in theta and alpha across two conditions (repeated and not-repeated) reveal attentional facilitation of certain items, which lead to conflict in probe matching for those items which are not facilitated during memory retrieval and can be associated with underlying causes for task-specific behavioral differences elicited by the participants.

## 2. Materials and Methods

### 2.1 Participants

Twenty-five participants (12 females; M(age) = 25.04 years, SD = 2.52 years, range: 21-32 years) were recruited for the study. All participants had a university degree or higher, were right-handed,-reported normal or corrected-to-normal vision, and declared no history of neurological or psychiatric disorders. Two participants’ data were not included in the analyses as one did not follow the proper instructions, and the other performed below the chance level in behavioral analysis of accuracy. Following this, data from a total of 23 participants (11 females; M(age) = 24.82 years, SD =2.12 years, range: 21-28 years) were included in the present study for further analysis.

### 2.2 Ethics statement

The study was carried out following the ethical guidelines and prior approval of the Institutional Human Ethics Committee (IHEC). Written informed consent was obtained from all participants before the commencement of the experiment, and they were remunerated for the time of their participation.

### 2.3 Stimuli and Trials

The working memory task for the study was designed and presented using Presentation® software (Version 23.0, Neurobehavioral Systems, Inc., Berkeley, CA) and displayed on a 22-inch LED monitor screen (60 Hz; 1920×1080 pixels) at a viewing distance of approximately 75 cm.

In both the main experiment and practice trials, visual working memory was tested using a probe-matching task **(Figure 1).** Participants were subjected to 280 trials in 6 blocks, with each block lasting 7-8 minutes and a break of around two minutes. Participants responded in a two-alternate forced-choice (2-AFC) manner using the left and right arrow keys of the keyboard for “No” and “Yes,” respectively.

**Figure 1.**
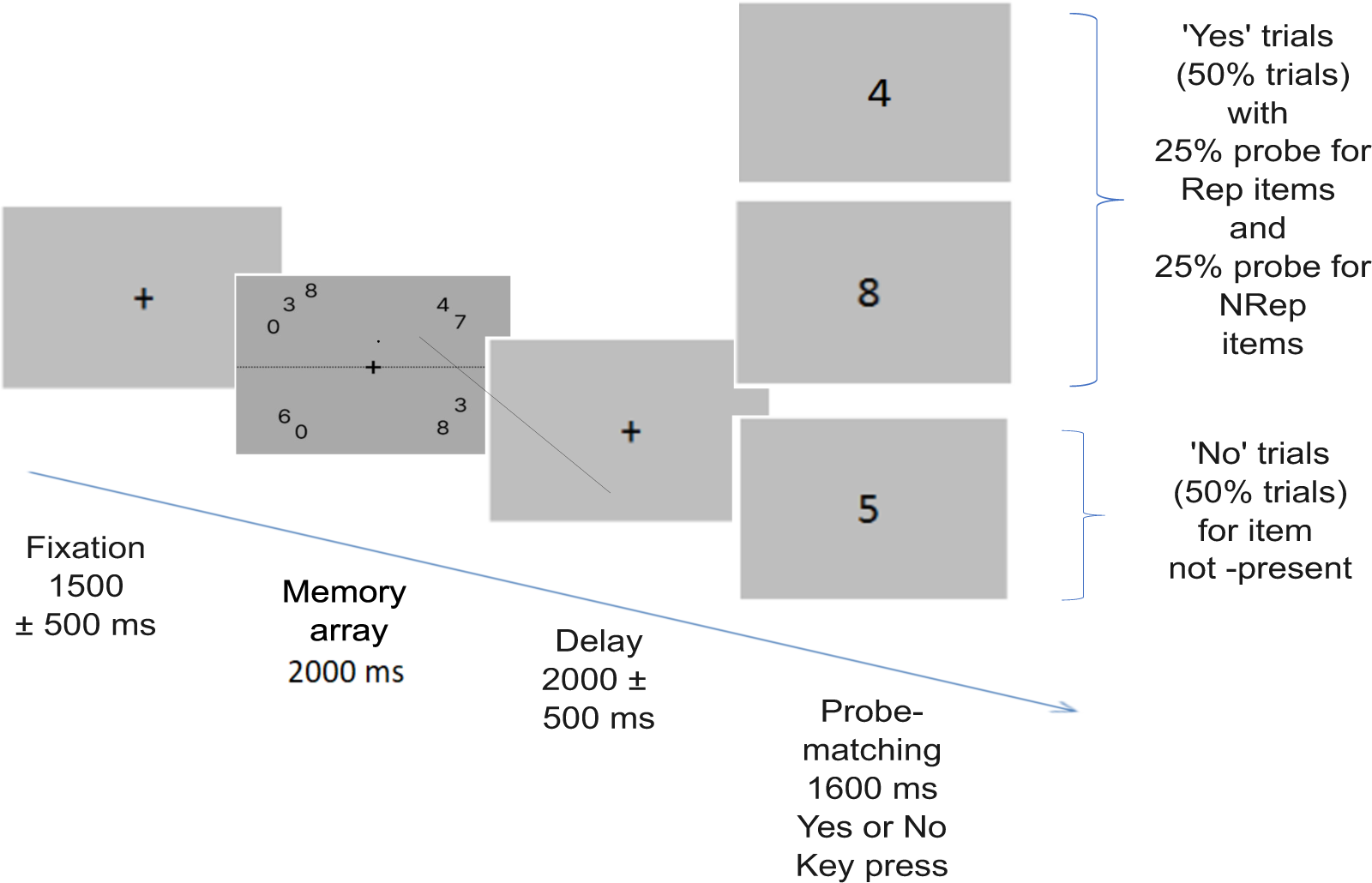
Trial structure for the probe matching task. Each trial begins with the presentation of one of the pseudorandomized memory arrays comprising of total 9 digits out of which three are repeated twice while remaining three are not repeated. After this a delay period occurs, followed by probe matching task. 50 % times probe matches item in memory array, with equal number of trials with probe of repeated items (Rep) and not repeated items (NRep) while 50 % times probe is for item not present in the memory array.

#### 2.3.1 Memory array

The memory array comprises of stimuli set where nine number digits were shown in each memory array, out of which three items were repeated twice and arranged in a jumbled fashion, along with three items that were not repeated; in total, nine items were presented in each memory array arranged in a circular fashion around fixation cross. These nine items are displayed with two to three items randomly shuffled between each quadrant to avoid encoding bias. Furthermore, sets of numbers used in a particular array were controlled to prevent the formation of commonly used chunks (e.g., numerical order, odd or even set, etc.). Numbers were shown within the foveal area (dva < 2.5 degrees), and each item subtends an angle of 0.76 degrees. Hence, all memory arrays were presented as stimulus images during the trials and were pseudo-randomized. Memory arrays were shown for 2000 msec on average.

#### 2.3.2 Trial structure

After presenting a black fixation cross for 1500 ± 500 msec, a memory array appeared for 2000 msec, which participants were instructed to remember. After the presentation of the memory array, a delay screen appears for 2000 ± 500 msec with a cross in the centre, followed by the onset of the probe on which participants had to respond whether the probe item was present in the memory array or not by using the left and right arrow keys. A black fixation cross was presented on a dark grey background throughout the trial. Participants were instructed to give responses as fast and accurately as possible. Response time and accuracy were estimated from behavioural data. The inter-trial interval (ITI) appears as a black screen after a response window of 1600 msec. Out of the total 280 trials, 140 were ‘No’ trials in which participants had to respond to a probe for which item was not present in the memory array. The remaining 140 ‘Yes’ trials were for probes having a corresponding item in the memory arrays. Out of all the ‘Yes’ trials, half had a probe for Rep items, and the other half had a probe for NRep items.

### 2.4 Behavioural analysis

Next, the response time and accuracy in the memory task were quantified for each participant. For response time and accuracy analysis, we used data from all the “Yes” trials of the Repeated (Rep) and Not-Repeated (NRep) categories, where response time and accuracy were calculated for the response window starting from probe onset till button press for Yes or No for probe matching. Data from two subjects were not included in the analysis, as one subject had poor accuracy (38%), whereas the other participant did not follow the instructions well. Outlier trials were removed using the Inter-Quartile range (IQR) method, where any data point less than 1.5 times the IQR below the quartile (Q1) or greater than 1.5 times the IQR above the quartile (Q3) is removed. Only trials with correct responses for probe matching in Rep and NRep conditions were used for response time analysis. After removing the outlier trials, response accuracy was analyzed. Two-tailed Wilcoxon signed-rank test was used to compare for significant differences in response time and response accuracy for Rep and NRep conditions. Effect size was quantified using *r* (the value of the z-statistic returned by the test, divided by the square root of the sample size).

## Data Acquisition and Analysis

### 2.5 EEG data

EEG recordings were obtained from 64 Ag/AgCl active electrodes (Brain Products GmbH, Gilching, Germany) using a Brain Vision Recorder. The 64-channel EEG signals were recorded using the International 10% electrode placement system and checked before and after the experiment. Reference electrodes were Cz, grounded to AFz. Channel impedances were kept at < 25 kΩ. Data were acquired continuously in AC mode (sampling rate, 1 kHz).

### 2.6 Pre-processing for EEG Signals

Analysis was conducted on twenty-two participants’ EEG data using MATLAB® and the EEGLAB toolbox (Delorme & Makeig, 2004). Data of one participant was discarded at this step due to very noisy recordings (with asymmetric variations of very large amplitudes due to a skull implant that the participant informed later). EEG data were down-sampled to 256 Hz, and High-pass (0.5 Hz) and low-pass filters (45 Hz, respectively) were applied before the data were re-referenced to the linked mastoid (TP9 and TP10). Noisy channels were removed after visualization of spectral power over those channels and removal of bad temporal segments. Next, we applied the Infomax independent component analysis (ICA) algorithm to detect artefactual ICAs (eye blinks, ocular, muscular, and electrocardiograph artefacts), and subsequently, these components were removed manually after visual inspection. Epochs of 0 msec to 1600 msec were extracted from the probe display onset till the end of the response window. They were sorted for the Rep and NRep two probe conditions and used for the Event-related spectral perturbation (ERSP) analysis. ERSP was computed using the *newtimef* function of the EEGLAB toolbox. The data was decomposed in a time-frequency domain across a frequency range from 3 Hz to 30 Hz using a complex Morlet wavelet. The pre-probe duration (-1000 to 0ms) was used as a baseline for baseline subtraction.

### 2.7 Spectral Analysis

The pre-processed EEG data was decomposed in a time-frequency domain across a frequency range from 3Hz to 30Hz and is computed by convolving three-cycle complex Morlet wavelets. These analyses were based on 200-time points from −1000 to 1600 msec, centered on the appearance of the probe till the end of the response time window, for the epoch corresponding to probe absence /presence, the number of cycles in the wavelet increased linearly from 3 Hz at the lowest frequency to 25.6 Hz at the highest frequency. The wavelet used to measure the amount and phase of the data in each successive, overlapping time window begin with a 3-cycle wavelet (with a Hanning-tapered window applied) and ‘0.8’ is the number of cycles in the wavelets used for higher frequencies will continue to expand slowly, reaching 20% (1 minus 0.8) of the number of cycles in the equivalent FFT window at its highest frequency. Baseline correction was applied by subtracting the mean power in the time window before the presentation of the probe (-1000 to 0 msec) from the power in the post-probe onset window. The ERSP was obtained by averaging the normalized representations across epochs, separately for the two probe conditions Rep and NRep. Epochs were baseline corrected by removing the temporal mean of the EEG signal on an epoch-by-epoch basis. Only trials where participants responded correctly to the probe were included in these analyses to observe the difference between probe matching for Rep and NRep categories.

### 2.8 Statistical Analyses of ERSP Power Changes

In the time-frequency analysis, we used the cluster-based permutation statistics to test for statistically significant differences in ERSP power for two probe-matching conditions (Rep and NRep) using EEGLAB’s toolbox’s *statcond* function along with false discovery rate (FDR) correction for multiple comparisons to estimate time-frequency clusters that were significantly different (with p < 0.05) between Rep and NRep conditions for the period around probe appearance till the end of the trial at 1600 msec Here, the null distribution is created by repeatedly shuffling the condition labels and recalculating the test statistic under the assumption of no true difference between conditions. The null distribution generated was used to compare the observed data clusters, which were considered significant if their difference exceeded the 95th percentile of the null distribution (p < 0.05, two-tailed). In addition, we visualized scalp maps for which ERSP values were averaged for different frequency bands for Rep and NRep conditions. The time period and frequency range for sensor analysis were decided based on identified significant clusters using time-frequency analyses and generated plots. We used cluster-based permutation statistics with 2000 iterations to identify sensors with statistically significant differences in ERSP power for the two conditions for all of our analyses using *statcond* function in the EEGLAB toolbox. Two-tailed paired t-tests with a threshold of 0.05 were used to identify significant clusters, along with permutation statistics, and to evaluate the sensors exhibiting statistically significant differences in ERSP power. Statistically significant clusters exceeded the 95th percentile of this null distribution. The details of all the specific individual analyses are further elaborated in the results section. Statistical analysis was carried out separately for alpha (8–12 Hz), theta (4-7 Hz), and beta (13-30 Hz) frequency ranges.

### 2.9 Data and Code Accessibility

All the behavioral and EEG data acquired from the participants and the analysis carried out during this study are available from the corresponding authors upon reasonable request. The pre-processed EEG data and codes/scripts used for all the analyses conducted in this paper will be made freely available to download from https://github.com/dynamicdip/

## 3. Results

### 3.1 Behavioural response

We only used data for trials with correct responses for probe matching in Rep and NRep conditions in the response time analysis. The violin plots **(Figure 2A)** (generated using ggplot2 (Wickham, 2011) in R software) for both conditions depict that the response times for probe matching follow Rep < NRep. The Response time distribution of the Rep condition is skewed and visually asymmetric. Hence, we employed a non-parametric two-tailed Wilcoxon signed-rank test to compute the statistical significance of differences between the medians of response times (RTs) of any two categories. Effect size was quantified using *r* (the value of the z-statistic returned by the test, divided by the square root of the sample size). We rejected the null hypothesis as we found using the Wilcoxon Signed-Rank test that there is a statistically significant difference between Rep and NRep with Rep having lower RT values (Median = 677.5, n = 25) than NRep (Median = 741.1, n = 25), (Z= 4.3589, p < 0.001, r = 0.87).

**Figure 2.**
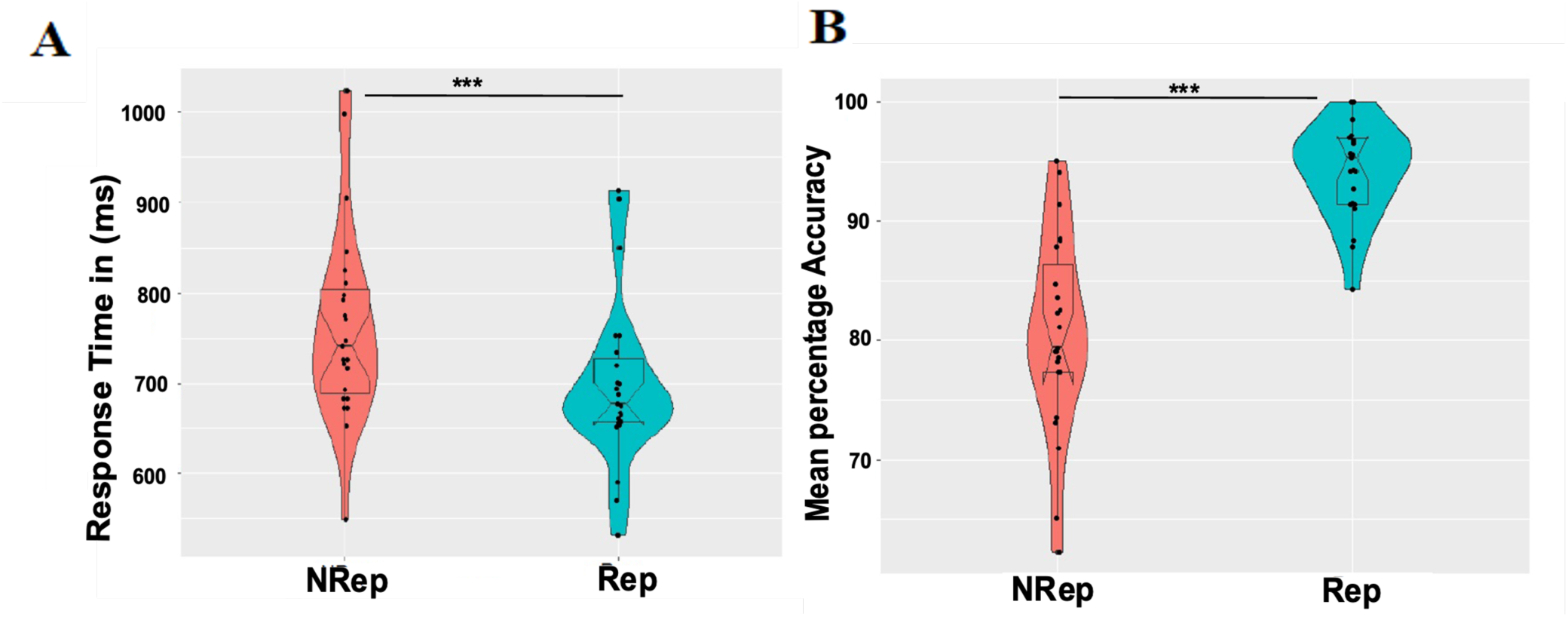
Behavioural Results. **A**, Each Violin plot shows responses time distribution for each condition (NRep Vs Rep). Each dot represents average response time for one participant. **Figure 2.B** shows Mean percentage accuracy distribution for each condition (NRep Vs Rep) where each dot represents average accuracy (in percentage) for one participant.

For response accuracy analysis, mean percentage accuracy (MPA) was calculated and plotted **(Figure 2B)** with distinguishable differences in distribution and median values of response accuracy using the two-tailed Wilcoxon Signed-Rank test. Effect size was quantified using *r* (the value of the z-statistic returned by the test, divided by the square root of the sample size). This Wilcoxon Signed-Rank test showed that response accuracy was significantly higher for the matching probe for Rep items (median = 100, n = 25) in comparison to that for NRep items (Median = 80, n = 25) with Z= -4.3589, p < 0.001, r = -0.87.

### 3.2 Event-related spectral perturbations in Rep versus NRep probe conditions

Next, we characterized whether the neural dynamics might reflect changes in spectral perturbations in different frequency bands due to these differences in response time and accuracy for the two probe-matching conditions.

In **Figure 3 (A)**, ERSP with data for both Rep and NRep conditions collapsed into one plot to visualize the grand average ERSP for values for frequencies ranging from 3 Hz to 30 Hz and for the temporal duration of -100 to 1100 msec post-probe presentation for ERSP plots, where 0 msec represents the onset of the probe across all subjects. This was done to avoid circularity in Time of interest (TOI) selection for analysis; instead, peak values of different EEG oscillatory rhythms are utilized based on data visualization. No statistical tests were performed at this level. Alpha was most prominently synchronized between 400 to 800 msec at 9-12 Hz. Also, event-related desynchronization was visible in the beta band between 13–21 Hz around the temporal window of 300 to 650 msec. The synchronization of the theta band is visible in the range of 4-7 Hz around the temporal window of 100-500 msec.

**Figure 3.**
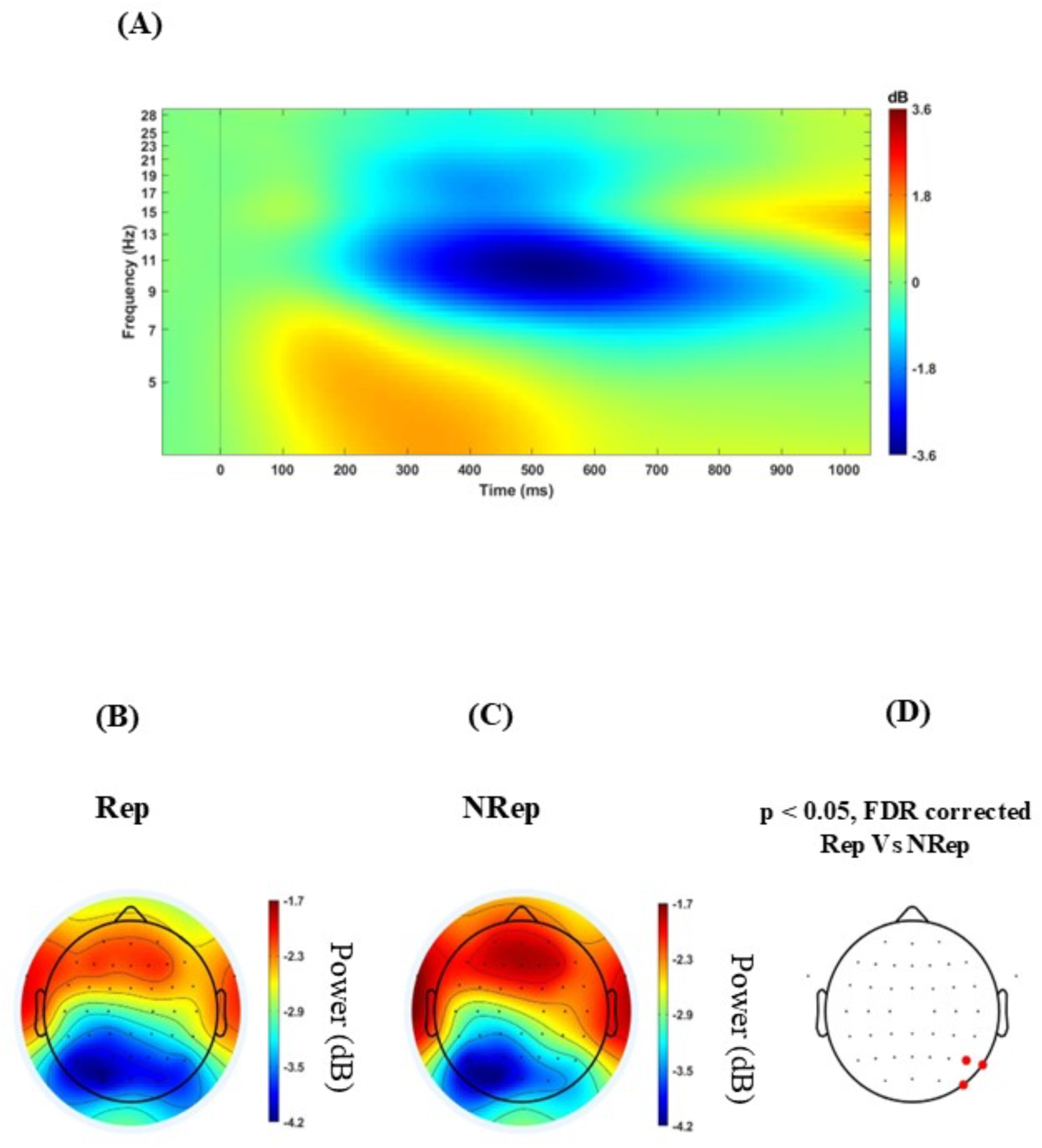
**A** Grand average of Event related spectral perturbation including trials of both the conditions across all the subjects in different EEG bands from 3 to 30 Hz. **Figure B, C** and **D** Scalp maps for all the electrodes averaged over frequency from 9 Hz to 13 Hz separately for each condition for 400 msec to 800 msec. Rep (left), NRep (middle), and **D** show a plot of FDR-corrected clusters with the threshold of 0.05 reflected on the vertical bar after cluster-based permutation showing a significant difference (red dotted) in PO8, P4 and P8 for ERSP-based topographies of two conditions for right parietal electrodes.

### 3.3 Topographical difference in Parieto-occipital Alpha power

Attention typically plays an important role in VWM retrieval, hence, we were interested in studying the role of the alpha band oscillations in mediating internal attention and suppressing irrelevant representation in WM during probe matching. Event-related alpha desynchronization was observed as depicted in **Figure 3B** and **C**. Furthermore, **Figure 3C** shows relatively increased alpha power in the right parieto-occipital area for NRep in comparison to Rep conditions as displayed in **Figure 3B**. The topoplots generated **(Figure 3B, C and D)** using cluster-based permutation statistics in the relevant temporal response window of 400 msec to 800 msec after probe onset and in the range of 9Hz to 12Hz frequency revealed significant involvement of parieto-occipital electrodes namely PO8, P4, and P8, showing enhanced power change in NRep compared to Rep conditions. We observed one negative cluster consisting of right parietal sensors, namely PO8, P4, and P8 [*t*_(21)_ = -2.97, p<0.001], with t-value peaking at -3.35 for PO8.

### 3.4. ERSP difference in Beta power for Rep vs. NRep probes

Next, we investigated beta band (13-20 Hz) desynchronization in C3, i.e., contralateral, which may be responsible for the response by the right hand with “Yes” for valid probe-matching, which indexes the prioritization of item in retrieval by enhancing the motor preparation for appropriate response selection in VWM for two conditions.

**Figure 4 A, B, and C** display ERSP plots for the C3 electrode averaged over frequency from 13 Hz to 20 Hz separately for each condition from -100 msec to 1100 msec around probe onset. Cluster-based permutation analysis revealed a significant (negative) cluster in beta band (13-20 Hz) between 200–400ms post-probe presentation over C3 electrodes with [t_(21)_ = -176.91, p = 0.018] peaking at 350ms and 16.5 Hz, showing stronger desynchronization of beta in Rep condition compared to NRep condition. NRep condition **(Fig 4 B)** indicates relatively delayed desynchronization of the beta band for the NRep compared to the Rep condition.

**Figure 4.**
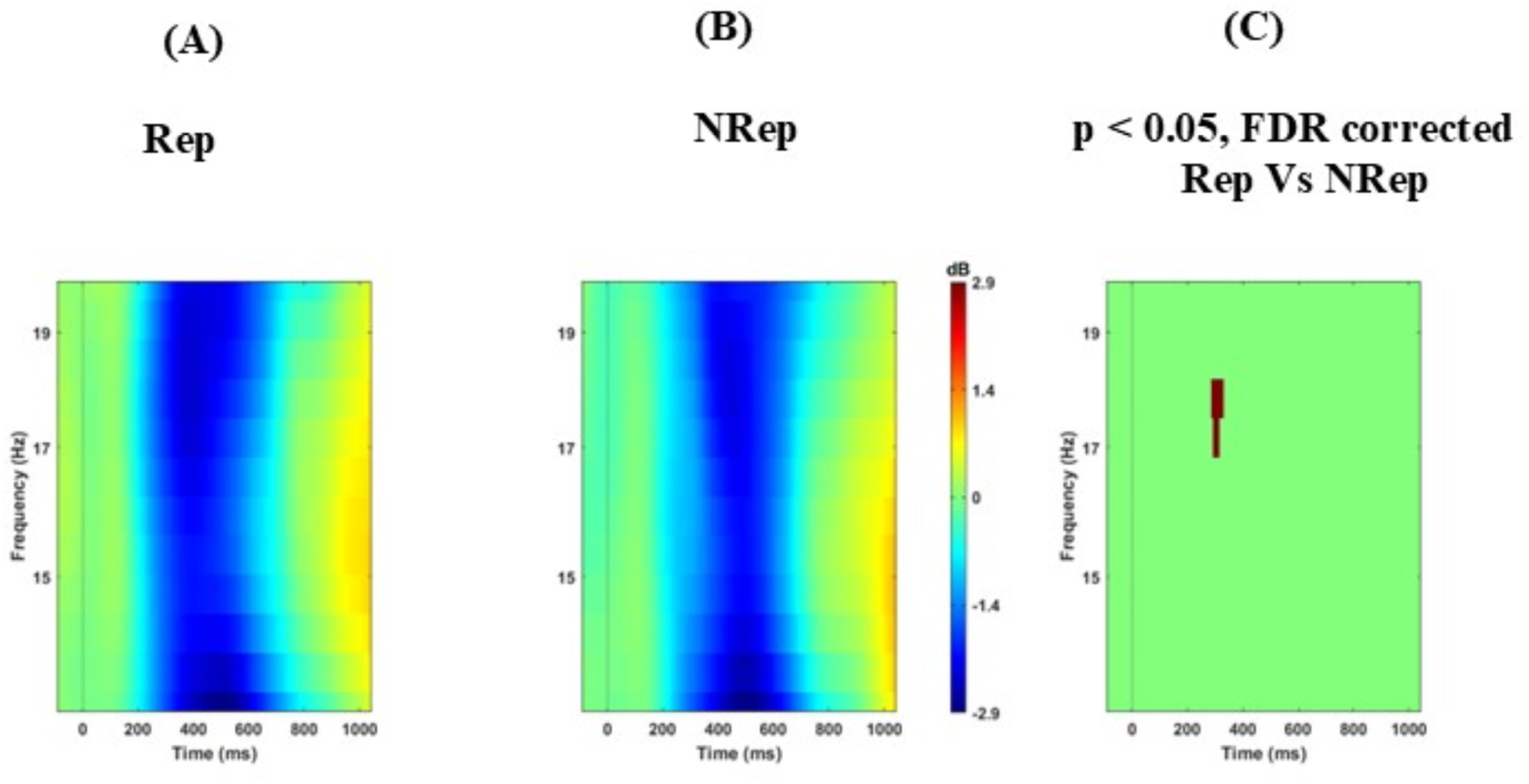
Beta power change for the two conditions. **A**, **B**, and **C** display ERSP plots for C3 electrode averaged over frequency from 13 Hz to 20 Hz separately for each condition from -100 msec to 1100 msec. Rep (left), NRep (middle), and plot of FDR corrected clusters (within white dashed lines) with the threshold of 0.05 after cluster-based permutation (right) showing significant difference in ERSP of two conditions. NRep condition shows positive cluster with delayed desynchronization of beta band here in comparison to Rep.

### 3.4 ERSP differences Frontal-medial Theta band oscillations

Next, we investigated the frontal-medial electrodes to examine the difference in theta power for the two probe conditions. Using cluster-based permutation, we found significant negative cluster at around 5 Hz - 7 Hz and 600 to 900 msec even after FDR correction with [*t*_(21)_ = -561.22, *p* = 0.022] peaking at 800 ms and 6.5 Hz, reflecting significantly higher theta power with a threshold of 0.05 for retrieving items using the probe for NRep category in ERSP plot **(Figure 5 A, B, and C).** The topographical distribution of ERSP **(Figure 5. D, E, and F)** over fronto-medial electrodes involving multiple sensors F1, F2, Fz, FCz, FC1, FC2, C1, and C2. The selection of these sensors was motivated by the previous studies (Ferreira et.al., 2019). In particular, C1 and C2 electrodes were used for analysis in place of Cz, as it was used as the reference electrode. The cluster of these sensors was depicted in **Figure 5. D, E, and F**, where permutation-based analysis was done for 5Hz to 7Hz and a period of 600 msec to 900 msec after probe onset. Multiple sensors, namely Fz, FC1, FC2, C1, C2, F1, and FCz, showed significant negative clusters in average ERSP power *with* t_(21)_ = -16.04, p <0.02, with t-value peaking at -2.66 for FC1, showing an increase in average ERSP power for the NRep condition over the Rep condition.

**Figure 5.**
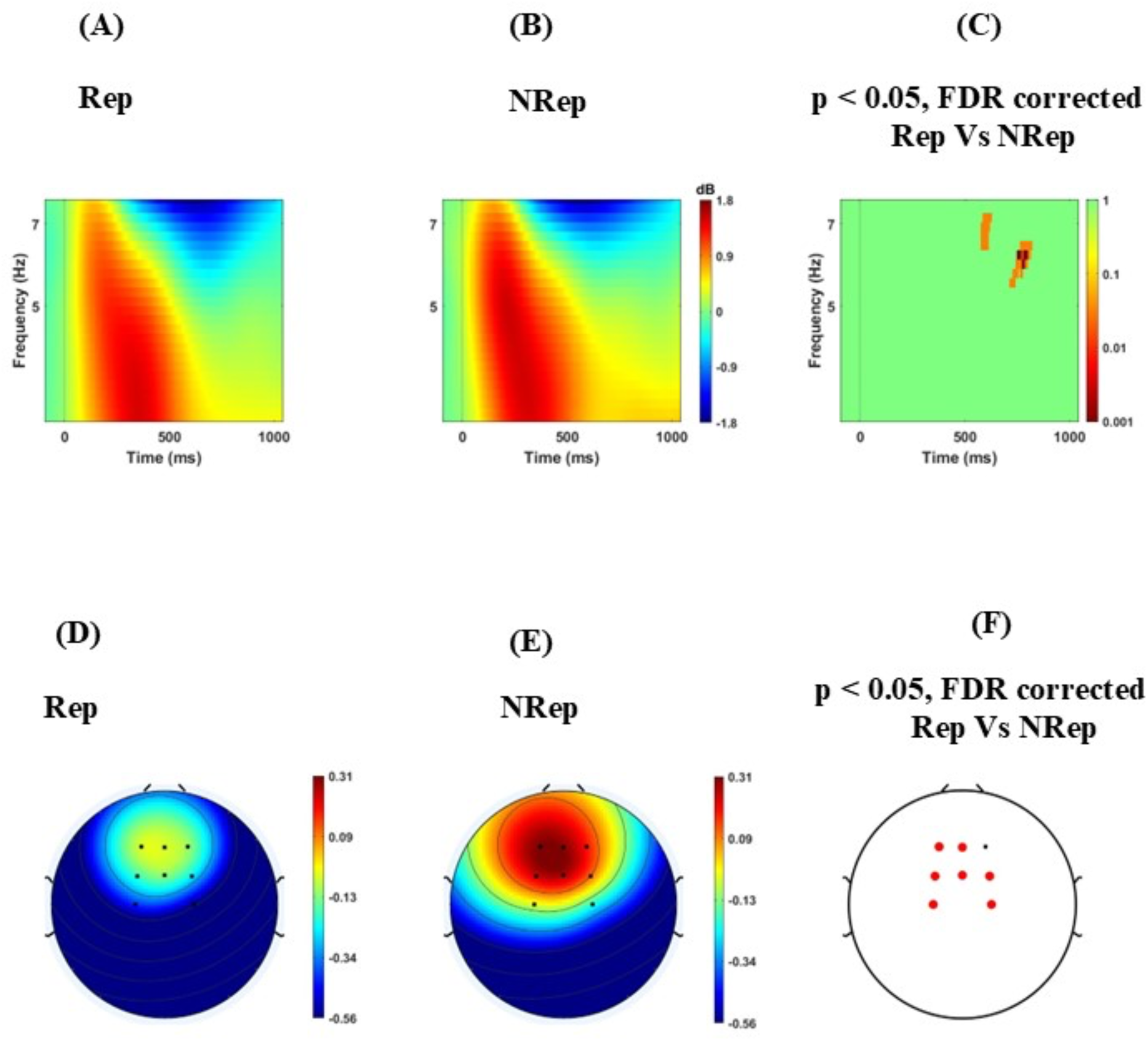
Increased Frontal–medial theta (FMT) power for responding for NRep items in comparison to Rep items. Figure **A**, **B** and **C** shows ERSP plots with positive clusters averaged over frequency from 4 Hz to 7 Hz separately for each condition for a period of -100 msec to 1100 msec. Rep (left), NRep (middle), and plot of FDR corrected clusters (white dash lines) with threshold of 0.05 reflected on the vertical bar after cluster-based permutation (right) showing significant difference in theta power ERSP of two conditions. Figure **D, E** and **F** Scalp maps for Fronto-medial electrodes averaged over frequency from 5 Hz to 7 Hz separately for each condition for 600 msec to 900 msec. Rep (Left), NRep (middle), and significant FDR corrected clusters in frontal region (red dotted) Fz, FC1, FC2, C1, C2, F1and FCz with the threshold of 0.05 after cluster-based permutation (right) showing significant difference in ERSP based topography of two conditions with increase theta power for NRep.

## 4. Discussion

The present study investigated the difference in probe comparison in a VWM task. Here, the difference in behaviour was empirically studied in terms of response time and accuracy for matching the relevant probe. Probes were matched with the maintained WM representations, where the probe acts as an attentional template to match with the relevant representation one at a time until its match is found. The most facilitated representations are in direct access, while other representations are brought into focus of attention sequentially. Subsequently, using ERSP analysis, we investigated how spectral perturbations of different brain oscillations differ in probe matching for the two probe conditions, Rep and NRep. Different brain oscillations provide further evidence for this bias in processing different items in the WM task.

Behavioural results showed that Rep items probes match faster and more accurately to relevant representations in comparison to the NRep probes. Our results provide evidence for the facilitation of Rep representations in probe matching, comparable to that of visual similarity in the working memory paradigm (Hamblin-Frohman et al., 2023), but by using the repetition of numbers as a linking feature between items (Oberauer et al., 2009). Our experimental results demonstrate that the default prioritization of Rep items is a feasible scenario in response selection, as they are facilitated when maintained and when they are retrieved for valid probe matching (Moorselaar et al., 2014).

Contrary to the bottom-up saliency view (Theeuwes, 1992; & Wolfe, 1994), which suggests that non-redundant dissimilar items should have gained prioritized access in VWM, as also seen in the Ranschburg effect, we found that repeated items were facilitated during VWM retrieval, reflecting an enhanced and stable representation of repeated items in VWM (Guofang Ren et al., 2023). One probable reason for such attentional facilitation is the chunking strategy for repeated items (Oberauer, Klaus 2019). This finding further implies that probe matching for repeated items requires less effort as their representation were in an active state for direct access. This further suggests the internal representation of attentionally prioritized Rep items might conflict with valid probe matching for NRep items, which require flexible allocation of attention to NRep items (Emrich et al., 2017).

Our ERSP results further revealed the role of different frequency bands in differential response selection, attentional demands, and conflict in decision making, which are required for probe matching of Rep and NRep probes in VWM. Beta power is mostly attributed to its role in sensory-motor function involving motor response selection, where it has been found to index the prioritization of items in VWM (Ding et al., 2024). Here, we predicted that the attentional template for Rep items’ representations is prioritized in the maintained WM, which facilitates WM recognition during valid probe matching. In this study, we find that the Beta band (13-20 Hz) in the C3 electrode is desynchronized at around 200ms shortly after probe presentation for repeated items and is significantly different in ERSP power for Rep vs. NRep conditions, which facilitates the right hand for valid probe matching and might be associated with faster and clearer motor preparation for response selection as also suggested by shorter response time and high accuracy for Rep over NRep in the behavioural results. Meanwhile, delayed beta desynchronization observed for the NRep condition suggests that when the attentional template tries to match NRep category, there is a conflict and delay for probe matching. The delayed prioritization compared to the Rep condition is probably due to the default prioritization of Rep items as seen in early desynchronization.

Parieto-occipital alpha has been shown to act as a marker for attention when selecting task-relevant information in the WM paradigms (Klimesch, W, 1999; Cavanagh & Frank, 2014). Active inhibition of non-relevant but distracting repeated item representations requires suppression during probe matching for not-repeated conditions. This is reflected in the increase in alpha power and comparatively weaker desynchronization for parieto-occipital electrodes when matching probe for NRep items compared to Rep shows active inhibition of irrelevant information as shown in previous findings (Benedek et.al., 2014; Erickson et.al., 2019).

Comparatively increased fronto-medial theta power for responding to probe matching for NRep items implies the role of cognitive effort for their selection and to resolve the conflict arising from matching the relevant probe. This is in line with conflict in WM retrieval and cognitive effort literature, (Jacobs et al., 2006; Zuure et al., 2020; Botvinick et al., 2001; Onton et al., 2005) where the change in power of the theta band resolves the conflict arising when the attentional template of a valid NRep probe matches with Rep representations. Increased theta power during NRep items probe matching appears due to interference by repeated item representations, similar to frontal-medial theta power effects of cognitive interference (modulated by distractor strength) as suggested by (Nigbur et al., 2011; Magosso et al., 2024). Previous research from (de Vries et al., 2018; Riddle et al., 2020) supports the crucial causal role of theta in prioritizing task-relevant information and potentially suppressing information that is no longer relevant for successfully guiding behavior. Contrary to that, maintenance and recognition of repeated items require lesser cognitive control and are facilitated by a chunking-like strategy. Theta power is comparatively lower due to the repetition enhancement-like effect, as the default prioritization of repeated items reduces the effort to retrieve. Here, we fixed the number of items so that varying working memory capacity does not affect the theta power. The cognitive control over WM items, as reflected in the frontal-medial theta power, also drives the alpha oscillations in sensors of posterior parietal cortices (Klimesch, W, 1999; Helfrich et al., 2017; Erickson et.al., 2019).

In summary, our study provides evidence for facilitation and prioritization of repeated items as seen in the behaviour for probe matching, where shorter response and higher accuracy for Rep items create conflict for processing passively maintained NRep items. This explains that items in VWM are maintained and retrieved in an order where prioritized representations are facilitated due to certain perceptual features like repetition, and followed by not-repeated items, which are less facilitated. The probe matching for NRep items showed delayed desynchronization of beta at the C3 electrode, which is characteristic of slow response preparation. The increase in parieto-occipital alpha power for NRep is due to active inhibition of irrelevant but default-prioritized Rep items during probe matching. FMT power increase suggests a link to resolving the conflict of matching NRep items over Rep items. This evidence provides an explanation for prioritization and facilitation of the inter-item feature of repetition interfering with the items that are not facilitated, even if they are relevant. Taken together, our study provides crucial empirical evidence of facilitation and prioritization of repeated items over non-repeated items and elucidates how different EEG rhythms might facilitate recognition of repeated items over goal-relevant, not-repeated items in VWM.

## Author Contributions

**Abhishek Singh Narvaria:** Conceptualization, Formal analysis, Methodology, Visualization, Writing – Original draft preparation. **Arpan Banerjee:** Funding acquisition, Project administration, Resources, Supervision, Validation, Writing – Review & editing. **Dipanjan Roy:** Conceptualization, Funding acquisition, Methodology, Project administration, Resources, Supervision, Validation, Writing – Review & editing.

## Acknowledgements

NBRC Core funds supported this study. DR was supported by SERB Core Research Grant (CRG) S/SERB/DPR/20230033 extramural grant from the Department of Science and Technology, Ministry of Science and Technology, Govt. India. DR and AB acknowledge the generous support of the NBRC Flagship program BT/MIDI/NBRC/Flagship/Program/2019: Comparative mapping of common mental disorders (CMD) over the lifespan.

